# Structure mapping of dengue and Zika viruses reveals new functional long-range interactions

**DOI:** 10.1101/381368

**Authors:** Roland G. Huber, Xin Ni Lim, Wy Ching Ng, Adelene Sim, Hui Xian Poh, Yang Shen, Su Ying Lim, Anna Karin Beatrice Sundstrom, Xuyang Sun, Jong Ghut Aw, Horng Khit Too, Peng Hee Boey, Andreas Wilm, Tanu Chawla, Ming Ju Choy, Lu Jiang, Paola Florez de Sessions, Xian Jun Loh, Sylvie Alonso, Martin Hibberd, Niranjan Nagarajan, Eng Eong Ooi, Peter J. Bond, October M. Sessions, Yue Wan

**Affiliations:** Bioinformatics Institute (A*STAR), 30 Biopolis Street #07-01, Matrix, 138671, Singapore; Stem Cell and Regenerative Biology, Genome Institute of Singapore, Singapore 138672, Singapore; Program in Emerging Infectious Diseases, Duke-NUS Graduate Medical School, 8 College Road, 169857, Singapore.; Computational Biology, Genome Institute of Singapore, Singapore 138672.; Department of Neurology, National Neuroscience Institute, 20 College Road, Singapore 169856, Singapore.; Stem Cell and Regenerative Biology, Genome Institute of Singapore, 60 Biopolis Street, Singapore 138672, Singapore; Department of Microbiology and Immunology, Yong Loo Lin School of Medicine, National University of Singapore, Singapore.; Immunology programme, Life Sciences Institute, National University of Singapore, Singapore.; Infectious Diseases, Genome Institute of Singapore, Singapore; Pathogen Molecular Biology, London School of Hygiene & Tropical Medicine, London, UK.; Institute of Materials Research and Engineering (IMRE), A*STAR, Singapore 138634; Department of Materials Science and Engineering, National University of Singapore.; GERMS platform, Genome Institute of Singapore, Singapore 138672.

## Abstract

Dengue and Zika are clinically important members of the Flaviviridae family that utilizes an 11kb positive strand RNA for genome regulation. While structures have been mapped primarily in the UTRs, much remains to be learnt about how the rest of the genome folds to enable function. Here, we performed secondary structure and pair-wise interaction mapping on four dengue serotypes and four Zika strains in their native virus particles and infected cells. Comparative analysis of SHAPE reactivities across serotypes nominated potentially functional regions that are highly structured, show structure conservation, and low synonymous mutation rates, including a structure associated with ribosome pausing. Pair-wise interaction mapping by SPLASH further reveals new pair-wise interactions, in addition to the known circularization sequence. 40% of pair-wise interactions form alternative structures, suggesting extensive structural heterogeneity. Analysis of shared pair-wise interactions between serotypes revealed macro-organization whereby interactions are preserved at their physical locations, beyond their sequence identities. In addition, structure mapping of virus genomes released in solution-as well as inside host cells-showed that other helicases, in addition to the ribosome, play a role in unwinding viral structures inside cells. Mutational experiments that disrupt in cell and in virion pair-wise interactions result in virus attenuation, demonstrating their importance during the virus life-cycle.

## Introduction

DENV and ZIKV viruses are members of the flavivirus genus of the Flaviviridae family of RNA viruses and are important human pathogens imposing a high economic and social burden worldwide^1^. DENV is predicted to infect 390 million people per year, resulting in DENV fever and in severe cases, death^1^. The ZIKV outbreak in Brazil was declared a public health emergency of international concern by the World Health Organisation in 2016 and has been associated with Guillain-Barre syndrome and microcephaly in infants^2^.

The genome of DENV and ZIKV viruses consist of a ~11kb long positive strand RNA that encodes a single polyprotein that is post-translationally cleaved into ten mature viral proteins, including three structural proteins (C, prM and E) and seven non-structural (NS) proteins (NA1, NS2A, NS2B, NS3, NS4A, NS4B and NS5)^3^. Beyond studying the primary sequence of such genomes, investigating how the genome is organized is important for understanding virus function^4^. Highly structured elements and long-range interactions in the 5’ and 3’ terminal regions, including the capsid region, of flaviviral genomes have been shown to be essential to both translation and replication of these viruses^5–8^. Additional local structures throughout the genome have been computationally predicted to exist by Proutski et al however, these predictions and their potential functional relevance to the viral life cycle have not been assessed^9^.

Here, we perform genome-wide RNA secondary structure and interactome mapping on all four serotypes of dengue (dengue 1-4) and four geographically distinct Zika viruses (African, Brazil, French Polynesia, and Singapore) – representing the known genetic diversity of this emergent virus. To circumvent limitations in RNA structure probing of virus RNAs *in vitro*, related to changes in solvent conditions, altered RNA-protein interactions and the absence of the virus envelope^10, 11^, we performed structure probing of dengue and Zika inside their native virus particles and in infected cells (**Figure 1a**). We observe that these genomes are indeed highly structured and identified new conserved structures across these viruses. Further, we show that many additional long-range interactions, besides the circularization signal, exist and are essential for the virus life cycle. Finally, comparison of in cell and in virion RNA interactions show that many interactions are disrupted in vivo, suggesting that they are being actively unwound by helicases in the cell.

**Figure 1.**
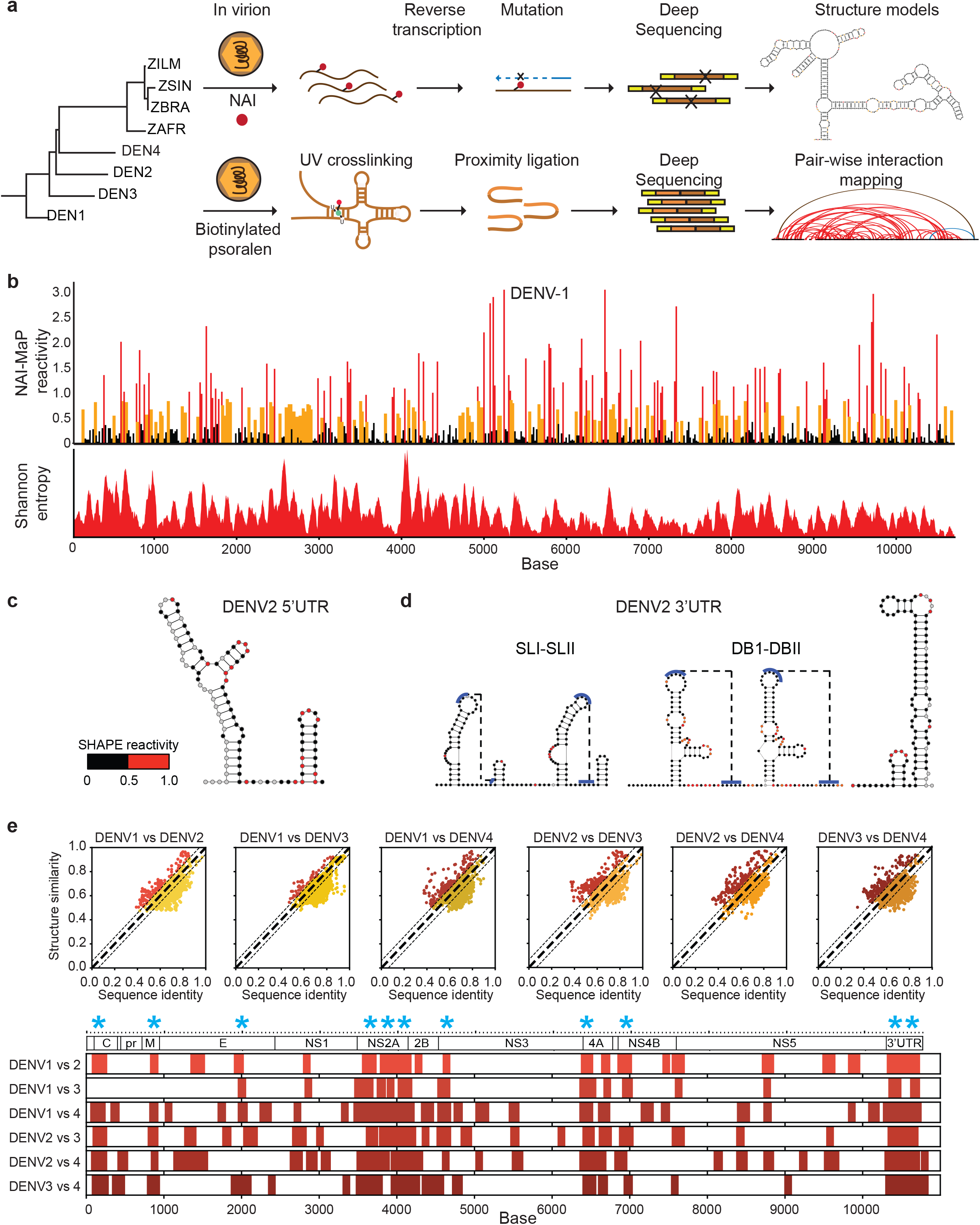
Genome-wide RNA structure mapping of dengue and Zika genomes inside virions. **a**, Schematic showing the workflow of identifying functional structural elements in eight viruses. Full length virus genomes are probed inside their virus particles using a SHAPE like chemical, NAI, which modifies single stranded regions along the genome. Pair-wise interactions within the virus genomes are also interrogated using biotinylated psoralen, which crosslinks base-paired regions inside the virus. The local and long-range experimental data are then used to constrain computational models to derive more accurate structure predictions for dengue and Zika. We also identify structurally conserved virus structures and determine the functional impacts of conserved and long-range interactions through mutagenesis and virus fitness assays. **b**, NAI-MaP reactivities and Shannon entropy along viral genome. Top, Raw NAI-MaP reactivities along dengue 1: Y-axis indicates the extent of NAI reactivity, and X-axis indicates position along the genome. Red, yellow and black bars indicates high, medium and low reactivities respectively. Bottom, Shannon entropy across the dengue 1 genome. Low values indicate higher probability of having a defined structure. **c,d**, NAI-MaP structure signals maps to known structures in the 5’UTR (**b**) and 3’UTR (**c**) of dengue1 virus genomes. The red circles indicate bases with high reactivity NAI-MaP signals, indicating that they are single stranded. **e**, Top: Scatterplot showing the sequence identity and structure similarity derived by NAI-MaP between each pair of dengue strains in 100nt windows. The red dots show locations whereby structure similarity is greater than sequence similar by 1 standard deviation from mean. Bottom: Each row indicates the locations of the high structure/sequence similarity regions (top) in each dengue pair, along the dengue genome. The structure conserved regions are consistently located at specific locations along the genome (200, 900, 2000, 3800, 4000, 4200, 4600, 6500, 7000, 10400 and 10600nt regions, starred), indicating that the high structure conservation is not a random and is an evolutionary constraint on structure beyond constraints on sequence alone.

## Results

### NAI-MaP of Dengue and Zika genomes inside virus particles

To map secondary structures across the viral genome, we treated intact virus particles with a SHAPE-like chemical, 2-methylnicotinic acid imidazolide (NAI)^12^, which has been shown to modify single-stranded regions along RNAs efficiently, *in vivo*. We then extracted the RNA genomes from the virus particles and identified the modification sites by mutational mapping (MaP) (**Figure 1a**)^12^. NAI-MaP signals have first been validated to be accurate *in vivo*, based on the 28S rRNA of Hela cells (**Extended data 1a**).

We performed at least two biological replicates of NAI-MaP on each of the four dengue and four Zika viruses, and sequenced >120 million reads per sample (**Supplementary Table 1**). This resulted in >100 reads mapped per base on 99.99% of all bases across the eight viruses, yielding structure information for most of the bases along each genome (**Supplementary Tables 2,3**). The NAI-MaP reads between two biological replicates were well correlated to each other (Pearson correlation, *p*>0.81), suggesting that the structure probing was of good quality. Gel electrophoresis of extracted dengue and Zika RNA showed that the majority of the genomic RNA is intact (**Extended data 2a,b**), indicating that structure probing was performed while the RNA was in its full-length context. NAI-MaP signals on renatured dengue 1 RNA are also largely congruent with the structure signals from Parallel Analysis of RNA Structures (PARS), which utilizes enzymatic footprinting coupled with deep sequencing^13^ (**Extended data 2c**).

### Dengue and Zika genomes contain extensive secondary structures in the coding regions

In addition to the known elements in the 5’ and 3’UTRs^6^, application of NAI-MaP to the dengue and Zika genomes showed that there are many previously unidentified structured regions across the entire genome (**Figure 1b, top**). We also calculated the Shannon entropies^12^, which indicates the likelihood that a region forms defined structures, along the entire length of dengue and Zika genomes to identify regions that are likely to form unique structures (**Figure 1b, bottom**). NAI-MaP reactivities mapped to known structures in the 5’ and 3’UTRs of dengue virus demonstrated that we could accurately detect single- and double-stranded regions within the dengue1 and dengue 2 genomes as expected (**Figure 1c,d, Extended data 2d**).

Structure probing across different viruses allows us to perform comparative structure analysis across the 8 viruses to identify shared structures. As the four dengue serotypes share around 60-70% sequence identities from each other and ~58% sequence identity with the Zika strains (**Extended data 3a**), the sequence divergence allow us to analyse the relationship between their structure conservation and sequence identity. For ease of comparing structural information across multiple genomes, we discretized the gradated NAI-MaP reactivities into a binary score that indicates whether a base is reactive (unstructured, reactivity >=0.5) or unreactive (structured, reactivity <0.5), based on the benchmark reactivity of the 28S rRNA (**Extended data 3b**). We observed that unpaired bases shared across the viruses tend to have higher sequence conservation (**Extended data 3c**), suggesting that these regions might contain important sequence information for gene regulation such as for interaction with RNA binding proteins. As expected, regions that share similar structure patterns between the 8 viruses generally share higher sequence identity, consistent with the idea that sequence plays an important role in RNA structure (**Figure 1e, Extended data 3d**). However, we also observed genomic regions that consistently show higher structural similarity over sequence similarity, such as in the NS2A region, suggesting that specific regions along the genomes are under evolutionary pressure for structure conservation (**Figure 1e, starred**).

To further identify potentially important structures along the genome, we searched for structurally similar, and also highly structured regions with low synonymous mutation rates. Synonymous mutation rates are calculated along hundreds/thousands of dengue sequences and hundreds of Zika sequences within each serotype, to identify mutation cold spots within the genome (**Extended data 4a,b, Methods**). We nominated sixteen dengue and twelve Zika consensus RNA regions that fulfilled two of three criteria, namely that they: (i) are highly structured; (ii) share similar NAI reactivity patterns; and (iii) show low mutation rates within and across serotypes (**Figure 2a, Extended data 4c, in purple, Methods**). The newly identified genomic locations have significantly lower Shannon entropies than average, indicating they are likely to form unique structures (**Extended data 4d**). They also overlap with the regions that show evolutionary pressure at the structural level (above), further confirming the importance of these regions.

**Figure 2.**
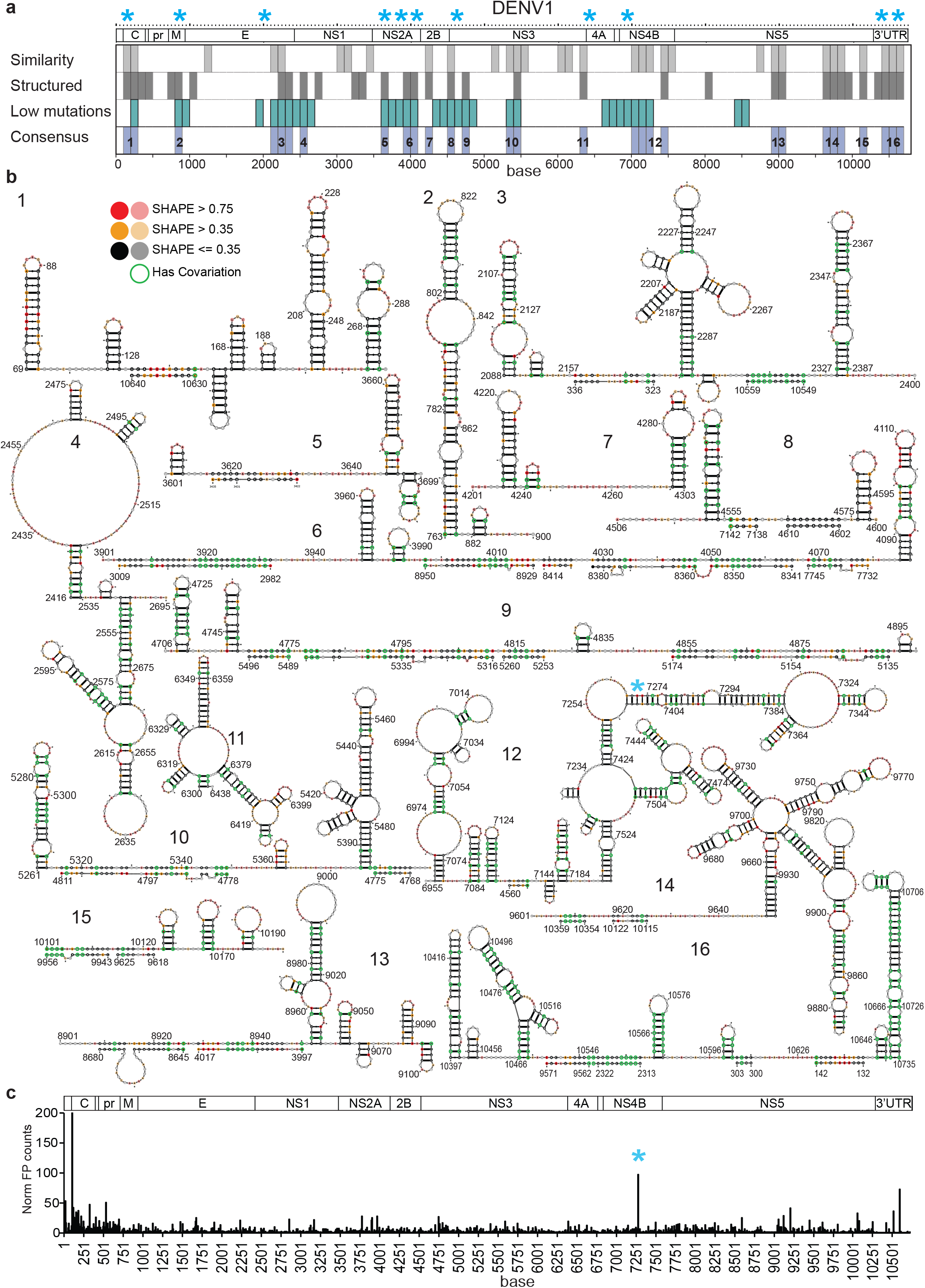
Dengue genome contains many conserved structural elements. **a**, Plots show 100 nucleotide regions across four dengue genomes that have highly similar structures (grey), are highly double-stranded (black), and accumulate low levels of synonymous mutations (blue). Regions that fulfil two out of three of the above criteria are selected as consensus regions (purple) and are potentially functionally important. The structurally conserved region from **Figure 1e** is starred in blue. **b**, Structure models of the dengue 1 genome using NAI-MaP as experimental constraints, for the 16 consensus regions identified in (**a**). The NAI-MaP reactivities (red, orange, black, indicating single, intermediate and double stranded regions respectively) and covariation information (green circles) are mapped onto the structures. **c**, Ribosome profiling data on dengue 1 in Huh7 cells. The Y-axis indicated the number of footprints/mRNA reads, and the X-axis indicated the position along the dengue1 genome. The strong ribosome pause site is starred in blue. The data is the average of 2 biological replicates of ribosome profiling.

SHAPE reactivities from structure probing experiments have been previously incorporated as pseudo-energies to generate accurate structure models for long RNAs using programs such as RNAstructure^14^. To model the RNA structures present within the dengue and Zika genomes, we incorporated our NAI-reactivities as pseudo-energies into the RNA structure program. We checked that incorporating NAI-MaP reactivities into structure modelling significantly improved the accuracy of the structure models using 16S and 23S rRNA (**Extended data 5a, Methods**). Structure modelling of the dengue and Zika genomes showed that it is highly structured, with numerous structures across the entire genome (**Figure 2b**). Predicted structures are significantly enriched for co-varied bases, confirming the high quality of the structure models (**Extended data 2b**). Interestingly, similar to other flaviviruses such as HCV^15^, all of the eight dengue and Zika genomes maintain a short median helix length of 4 consecutive canonical base pairs, with >90% of the base-pairs existing in helices that contains 7 consecutive base-pairs or less, enabling the genomes to evade immune surveillance inside cells (**Extended data 3c,d, e**).

To determine whether any of our structures in dengue and Zika could be associated with cellular processes such as translation, we performed ribosome profiling on human liver cells (Huh7) infected with dengue 1 virus^16^. Ribosome profiling showed the classic three-nucleotide periodicity of translating ribosomes on the genome (**Extended data 7a**), with a large pileup of footprints at the start of the dengue 1 polyprotein, suggesting that ribosome profiling data is of good quality (**Extended data 7b**). Globally, we did not observe a correlation between ribosome pausing and increased structure along the dengue genome (**Extended data 7c**), although we did observe a strong ribosome pause site in the middle of the coding region for the viral membrane protein NS4B (**Figure 2c**). Interestingly, the ribosome pause site resides in one of our dengue consensus structures that has low Shannon entropy (segment 12, **Figure 2b, Extended data 7c**). Structure modelling of this region indicates that the ribosome pause site is located near the base of a long stem, in a highly structured environment (**Figure 2b,c**, starred), suggesting that structure could play a role in translation pausing. To rule out that the strong ribosome pausing could be due to codon usage, we calculated the frequencies of codons around the ribosome pause site. We did not observe a significant difference in codon frequencies, as well as the presence of any rare codons, around the pause site, suggesting that codon usage is unlikely to be the main reason behind the strong ribosome pause (**Extended data 7d**). Further work needs to be done to understand the role of RNA structure in regulating translation in this region.

### Long-range pair-wise interactions are abundant in Dengue and Zika genomes

Beyond local RNA base pairing, long-range RNA-RNA interactions have been shown to play important roles in viral replication by enabling the genome to circularize and position the RNA-dependent RNA polymerase close to the transcription start site^5, 7, 17^. The three principal RNA sequences of the dengue genome responsible for the long-range interaction of the 5’ and 3’ ends are: (i) the circularization sequence (CS); (ii) the upstream AUG region (UAR); and (iii) the downstream AUG region (DAR)^17^. A fourth interaction region located at 150 bases (C1 structure of the capsid protein) and the dumbbell of 3’UTR has also been found to be involved in genome circularization^7^. To directly capture long-range pair-wise interactions that span longer than 500 bases in dengue and Zika genomes comprehensively, we performed two biological replicates of sequencing of psoralen crosslinked, ligated, and selected hybrids (SPLASH) on each of the eight viral genomes, inside their viral particles, using biotinylated psoralen, proximity ligation and deep sequencing (**Figure 1a**)^18^. We further enriched our data for real interactions by filtering against random interactions that could occur by permutation (**Extended Figure 8a**).

The resultant SPLASH data revealed thousands of new intramolecular long-range RNA interactions inside virus particles (median distance of interaction = 3.6kb), greatly expanding the list of long-range interactions known for any family of RNA viruses (**Supplementary Tables 4,5**). We observed interactions that arise from the known circularization sequences in both the dengue and Zika viruses as one of our top interactions (**Figure 3a**), indicating that genome circularization is not only important in vivo, but is also maintained in virions. To increase the resolution for identifying interaction positions, we performed peak calling of the mapped reads on each end of the interaction (**Methods**). Peak calling revealed clear peaks that were around 50 bases wide, centred at the known 5’ and 3’ UAR and DAR interaction sites (**Figure 3b**). Hybridization of the 5’ and 3’ peak regions using the program RNAcofold revealed the interaction sites of the UAR, DAR and CS regions in the dengue genomes (**Figure 3b**)^19^, demonstrating the precision of SPLASH data.

**Figure 3.**
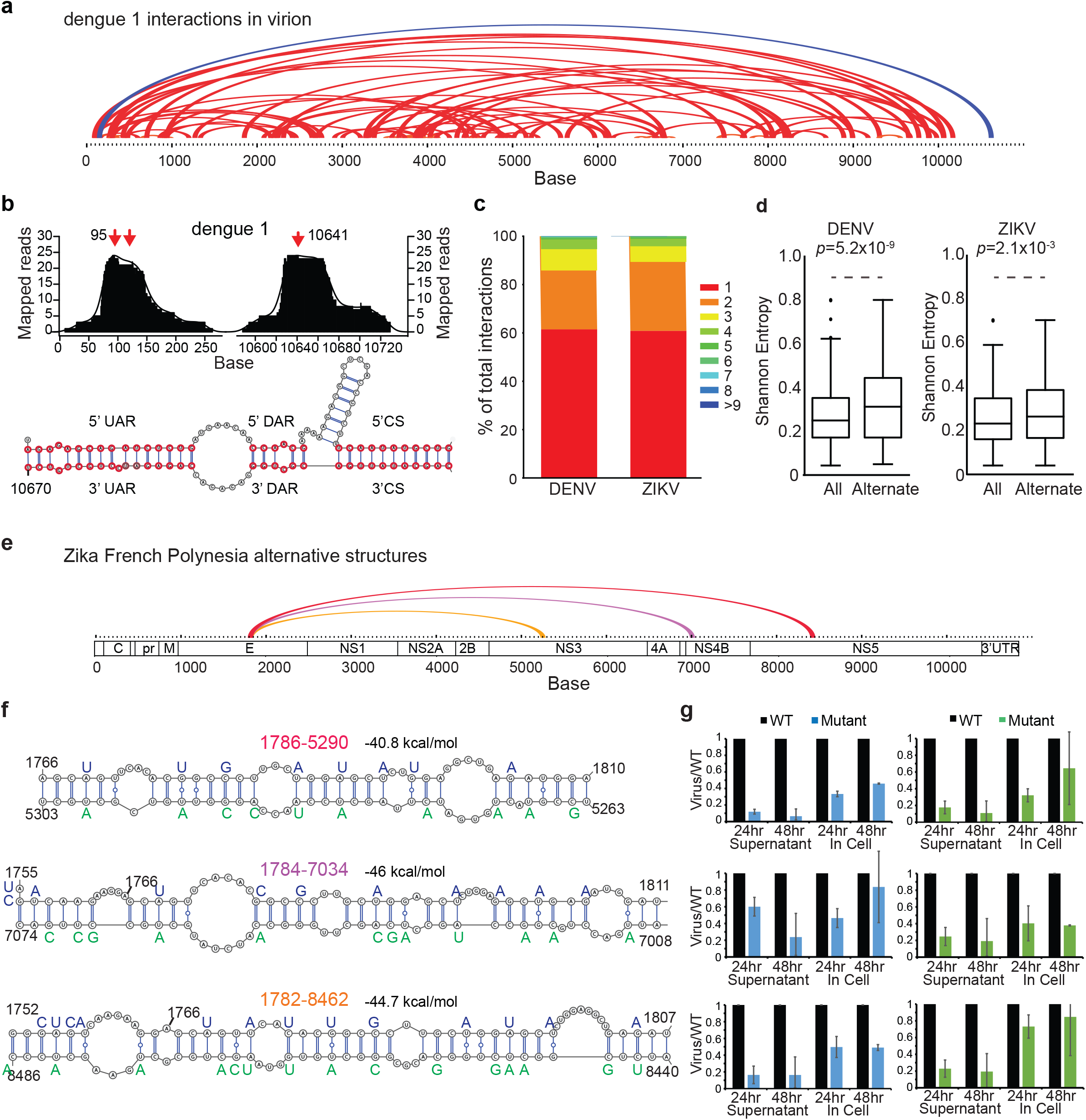
Dengue and Zika genomes forms extensive long-range interactions and forms alternate structures. **a**, Arc plot showing the most abundant top 75 pair-wise interactions in the dengue1 genome inside virions. The known circularization signal is coloured in dark blue. **b**, Top, Peak calling of the circularization interaction region shows that the peak of base-pairing occurs at positions 95 and 10641. Bottom, Prediction of hybridization using RNAcofold program using sequences around the called peaks identified three of the best known circularization signals. **c**, Bar charts showing the number of peaks that interact uniquely and with other regions along the genome. 40% of dengue and Zika interactions are not unique. **d**, Boxplot showing that regions with alternative base-pairing (alternate) have higher Shannon entropies than all regions present in the genome (all) for both dengue 1 and Zika French Polynesia strain. *P*-value calculated using T-test. **e**, Arc plot showing one set of alternative interactions found along the Zika French Polynesia genome. **f**, Structure models of the pair-wise interactions for the alternative structures in (**e**), using the program RNAcofold. Mutations along each strand of the pair-wise interaction are in blue and green respectively. **g**, qPCR analysis of the amount of mutant (from **f**) and WT virus that are produced in Huh7 cells and supernatant, 24 and 48 hours post infection. The Y-axis shows that amount of mutant virus produced as compared to wildtype.

### Dengue and Zika genomes forms extensive alternative structures

Previous studies in HCV suggests that its genome can fold into alternative conformations^20^. To determine the amount of structural heterogeneity in the dengue and Zika genomes, we calculated the number of RNA regions that are observed to interact with two or more other regions along the genome. We observed that ~40% of the genome folds into alternative structures, indicating a large amount of structural heterogeneity within the genomes (**Figure 3c, Extended data 8b**). Interestingly, the alternative interaction sites are enriched in regions with significantly higher Shannon energies, confirming that these regions tend to take on more than one structure (**Figure 3d**). To test whether the different alternative structures are generally functional or whether only the most stable interaction is important to the virus, we mutated long-range interactions that could pair with three different regions along the Zika French Polynesia genome (1786:5290, 1784:7034 and 1784:8462), disrupting the interactions that bring the nucleotide sequences encoding the envelop protein into close proximity with those of NS3, NS4B and NS5 respectively (**Figure 3e, Extended data 8c**). As we chose mutations that disrupt pair-wise interactions and preserve the amino acid sequence and codon frequencies, we were unable to design for compensatory mutations (**Figure 3f**). Mutations along each individual strand in the different alternative structures resulted in a decreased amount of virus produced in both the supernatant and inside infected Huh7 liver cells (**Figure 3g**). Interestingly, we observe a greater decrease in virus production in the supernatant as compared to inside cells, suggesting that the mutations might affect genome packaging.

### SPLASH reveals two distinct modes of conserved long-range interactions

Interactome mapping across the four dengue and four Zika strains allow us to identify shared long-range interactions that are present across different viruses. In addition to the known circularization signal, we also observed macro-patterns of conservation whereby genomic sequences from similar spatial locations consistently interact with each other, in two or more different viruses (**Figure 4a,b, Supplementary Tables 6,7**). We observed two distinct types of conserved pair-wise interactions. The first type preserved both the sequence and the spatial locations of the interactions across serotypes. This is exemplified by the discovery of a conserved long-range interaction between E and NS4B (bases 1200:7000) in dengue and Zika - whereby different viruses use homologous sequences to form the long-range interaction (**Figure 4c**). The second type of interaction preserves the spatial location, but not the sequence of interaction between the serotypes, to bring two distant regions of the genome into close proximity. This latter type of interaction is exemplified in the pairings between C and NS5 and NS2A and NS5 regions in both dengue and Zika genomes (**Figure 4d, Extended data 9**). While dengue 1 uses bases 218-253 (within capsid) to pair with bases 9899-9935 (within NS5), dengue 3 uses a similar region – bases 237-274 to pair with a different downstream region of 9800-9840 to form a stable structure. In Zika viruses, bases 351-369 are used to hybridize with bases 9629-9648 instead. This second mechanism of genome interaction, whereby different sequences could be utilized for genome organization, could hypothetically enable RNA viruses to tolerate mutations - as long as the overall architecture of the genome is not affected - and could also reflect convergent evolution in different viruses to achieve similar genomic folds.

**Figure 4.**
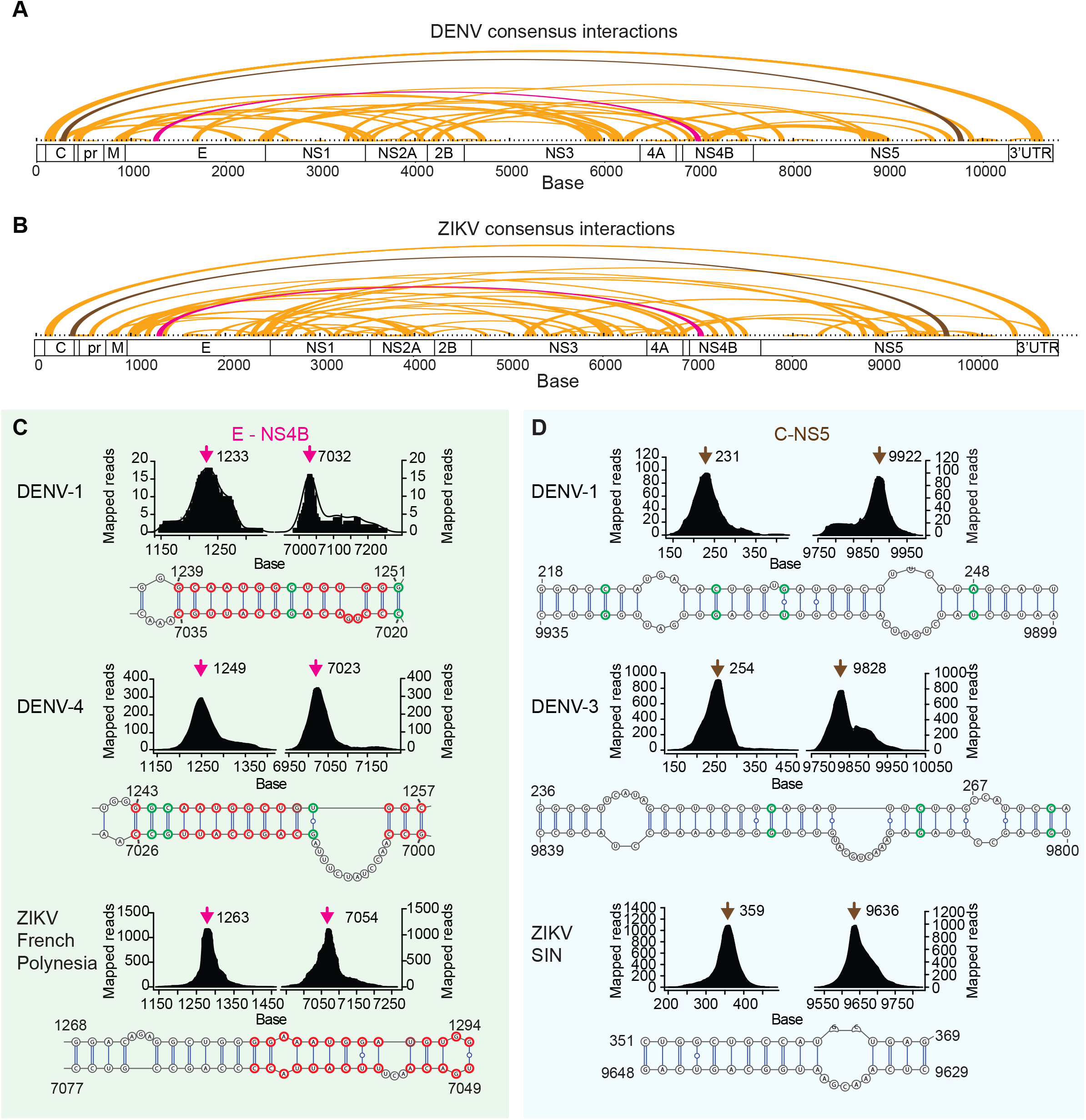
Two different modes of long-range interactions in dengue and Zika. **a,b**, Arc plots showing the top 100 conserved long-range interactions that are present in two of more different dengue (**a**) and Zika viruses (**b**). The pink and grey interactions are further characterized in **c,d. c**, New conserved long range interaction in dengue and Zika viruses that use similar sequences to bring E and NS4B regions together. Top, SPLASH interaction peaks between E and NS4B for different viruses. Bottom, Predicted hybridization models for the long-range interaction using RNAcofold. Homologous and co-varied bases between the different viruses are circled in red and green respectively. The pink arrows indicate the position of the peak of the interaction. **d**, New long-range interaction in dengue and Zika viruses that use different sequences to bring NS4 and NS5 regions close together. Top, SPLASH interaction peaks between NS4 and NS5 for different viruses. Bottom, Predicted hybridization models for the long-range interaction using RNAcofold. The grey arrows indicate the position of the peak of the interaction.

### Longer-range interactions are actively disrupted inside cells

While structure probing of virus genomes inside their virions provides useful information on their packaging, virus genomes exist in very different states inside their host cells. To understand how dengue and Zika genomes fold inside infected cells, we performed pair-wise interactome mapping of dengue1 and the four Zika viruses in human liver and neuronal precursor cells respectively (**Extended data 10a, Supplementary Tables 8,9**). We observed that the circularization signal is present inside cells, agreeing with its importance in genome replication^5^ (**Figure 5a, Extended data 10b**). Interestingly, we observed that the viruses form much shorter pair-wise interactions inside cells as compared to inside virion particles (**Figure 5b**), suggesting that the genomes are less structured inside cells. We hypothesized that the genomes are either more structured inside virions because of the spatial constraints induced by the virus envelop, and/or that the genomes are actively being unwound in vivo. To test this, we performed interactome mapping on virus genomes that are released from the virions by performing proteinase K treatment on the virus particles (**Figure 5c**). Although we do observe genome rearrangements when the genome is released in solution, in particular longer-range interactions are disrupted while new shorter-range interactions are formed, the genome remains highly structured in solution (**Figure 5c,d**). This suggests that the unstructured state we observe inside cells is a result of active unwinding by enzymes in vivo^21^.

**Figure 5.**
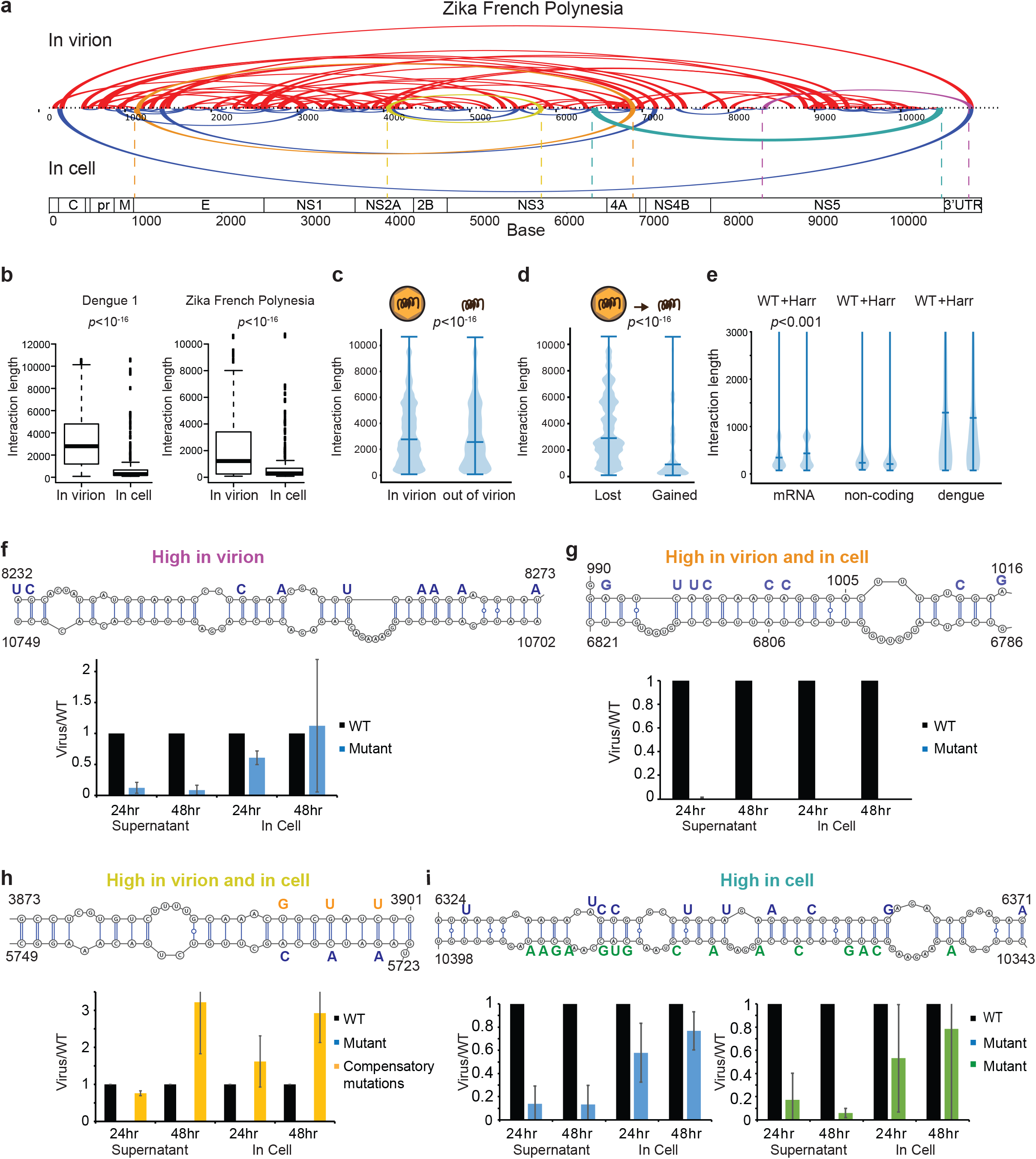
Long-range interactions in virions are disrupted inside host cells. **a**, Arc plots showing long-range interactions in Zika French Polynesia strain that are present inside virion particles (Top) versus inside host cells (bottom). **b**, Boxplot showing the distribution of pair-wise interaction lengths inside virions versus inside host cells for dengue 1 and Zika French Polynesia strain. Long-range interactions are significantly disrupted inside host cells. *P*-value is calculated using Wilcoxon Ranked Sum test. **c**, Boxplot showing the distribution of pair-wise interaction lengths inside virions versus when released into solution by proteinase K treatment. While there is a slight decrease in longer-range interactions, the dengue genome remains highly structured in solution. **d**, Boxplot showing the length of pair-wise interactions that are disrupted versus new interactions that are formed when the dengue genome is released into solution. Longer-range interactions tend to be disrupted while shorter-range interactions tend to be formed upon genome release. **e**, Boxplot showing the distribution of pair-wise interaction lengths of human mRNAs, non-coding RNAs and dengue1 with and without harringtonine treatment, inside Huh7 cells. mRNAs show increased structure upon translation inhibition, but not non-coding RNAs or dengue1. **f**, Top, Mutations (blue) are performed along a strand of the pair-wise interaction that is found dominantly in virions, as compared to in cell. Bottom, qPCR analysis of the amount of mutant and WT virus that are produced in Huh7 cells and supernatant, 24 and 48 hours post infection. The Y-axis shows that amount of mutant virus produced as compared to wildtype. **g**, Top, Mutations (blue) are performed along a strand of the pair-wise interaction that is found in to be abundant in both virions and cells. Bottom, qPCR analysis of the amount of mutant and WT virus that are produced in Huh7 cells and supernatant, 24 and 48 hours post infection. The Y-axis shows that amount of mutant virus produced as compared to wildtype. **h**, Top, Mutations (blue) and compensatory mutations (yellow) are performed along a pair-wise interaction that is found in to be abundant in both virions and cells. Bottom, qPCR analysis of the amount of mutant, compensatory mutant, and WT virus, that are produced in Huh7 cells and supernatant, 24 and 48 hours post infection. The Y-axis shows that amount of mutant virus produced as compared to wildtype. **i**, Top, Mutations (blue, and green) are performed independently along each strand of a pair-wise interaction that is found to be abundant inside cells. Bottom, qPCR analysis of the amount of mutant and WT virus, along each strand, that are produced in Huh7 cells and supernatant, 24 and 48 hours post infection. The Y-axis shows that amount of mutant virus produced as compared to wildtype.

As the ribosome is a major helicase that disrupts RNA structures *in vivo* and previous reports have shown that translation preferentially disrupts longer-range interactions^22^, we further tested the effect of translation inhibition on structure by performing SPLASH on dengue infected cells that are treated with harringtonine to immobilize ribosomes right after initiation^23^. As expected, inhibiting translation partially restored long pair-wise RNA interactions in human mRNAs, and not in human non-coding RNAs^22^. Surprisingly, we observed a negligible effect on RNA pairing in the dengue genome with and without translation inhibition (**Figure 5e**), suggesting that unlike cellular mRNAs, translation is not the dominant activity that remodels RNA structures for the virus. This opens up the possibilities that other helicases, including the viral helicase that is important for genome replication, could play dominant roles in remodelling viral structures inside cells.

As many of the longer-range interactions in virions are disrupted inside cells, it is unclear whether these pair-wise interactions are functionally important. To determine the significance of the pair-wise interactions that are present *in virion, in cell*, or in both, we performed mutagenesis experiments to disrupt the interactions found in all three classes (**Figure 5f-i**). Disruption of structures that are present in high abundance either inside virion particles or inside cells resulted in virus attenuation. We observed less virus production in the supernatant, as compared to inside cells, for two out of four interactions, indicating that genome packaging is a major stage of virus lifecycle that the interactions are involved in (**Figure 5f,i**). Interestingly, mutating two interactions that abundantly present in both the cells and the virus particles resulted in severe phenotype of >100 fold decrease in virus production (**Figure 5g,h**). While it is extremely difficult to design compensatory mutations to rescue pair-wise interactions in the virus coding region, due to amino acid and codon frequency constraints, we managed to design point mutations to rescue one of the key interactions between NS2A and NS3. While mutations in NS3 region (5723-5750) resulted in a severe virus attenuation, this phenotype is restored by designing compensatory mutations in the NS2A region (3871-3901) (**Figure 5h**). These results confirm that the newly identified pair-wise interactions play important roles during the lifecycle of the viruses.

## Discussion

Elucidating the molecular underpinnings of RNA viruses is key to understanding their pathogenesis^4^. Here, we integrated high throughput secondary structure mapping, pair-wise interaction mapping, ribosome profiling and evolutionary conservation information for four dengue and four Zika viruses to study how the structural organization of the genome impacts virus function^16, 18^. Importantly, the structure mapping is performed inside native virus environments, including inside virion particles and host cells, enabling us to identify biologically relevant structures. We discovered (i) highly structured and conserved elements across the dengue and Zika genomes, (ii) a new structural element associated with strong ribosomal pausing in the coding region of NS4B, (iii) alternative and conserved pair-wise interactions across serotypes, and (iv) that longer-range pair-wise interactions are actively disrupted inside host cells. Many of the shared pair-wise interactions preserve sequence locations but not sequence identities, suggesting an alternative mechanism by which viruses could use diverse sequences to achieve the overall goal of genome packaging. Surprisingly, while the ribosome has been shown to be a major helicase to unwind RNA structures during translation^21, 22^, we did not see a similar effect on dengue structure upon translation inhibition, as compared to host cells. This suggests that unlike mRNAs, there are likely other helicases involve in genome replication that cause structure remodelling of dengue inside cells.

Our mutagenesis experiments on pair-wise RNA interactions dominantly present inside virion particles, or inside cells, or both, showed many of the new interactions are likely to be functional during different stages of the virus life-cycle. Interestingly, we observe a more severe mutation phenotype for interactions that are present in both the virions and in cell, suggesting that these interactions are likely to be highly important across different virus life stages. Many of our structure disruptions appear to affect the amount of virus particles released into the supernatant more than inside cells, suggesting that they may impact genome packaging. This agrees with previous literature on HCV showing disruptions of genome interactions can affect genome packaging^24^.

In summary, our data expands upon the known structures in dengue and Zika and reveals a network of new interactions within the dengue and Zika genomes that are likely work together to support and facilitate virus fitness. This comprehensive resource of dengue and Zika genome organization provides the first foray into higher order genome organization of large RNA viruses and aids in the design of broad based RNA therapeutics in targeting the structure of these pathogenic viruses.

## Methods

### NAI-MaP structure probing inside virions

Dengue virus serotypes 1-4, belonging to EDEN viruses 2402, 3295, 863 and 2270 respectively (Genbank ID: EU081230.1, EU081177.1, EU081190.1, GQ398256.1), were amplified in Vero cells and harvested from the cellular media 4-5 days post-infection. Zika virus African strain (AY632535_MR766), French Polynesia strain (KJ776791_HPF2013), Brazil strain (KU365780_BeH815744) and the Singapore strain (KX813683_ZKA-16-097) were amplified in C6/36 cells and harvested 3-5 days post-infection.

Freshly collected viruses were centrifuged at 14000 rpm for 10 min at 4°C to remove cellular debris. The viruses are then separated into 3 reactions: 1) we added 1:20 the volume of 1M NAI (03-310, Merck, 25 μl of NAI in 500 μl of virus in cell media) and incubated the reaction for 15 min at 37°C for structure probing; 2) we added 1:20 volume of DMSO to the virus and incubated the reaction for 15 min at 37°C as negative control; 3) we set aside a third portion of the virus as for denaturing control in the downstream library preparation process. We then extracted all the viruses using TRIzol LS reagent (Thermo Fisher Scientific) following manufacturer’s instructions.

We performed gel electrophoresis of the virus RNA after each extraction on a 0.6% agarose gel, using ssRNA (NEB) as the ladder, to ensure that the virus RNA is intact. We typically see a single band above the 9kb RNA ladder that indicates the presence of intact full length dengue and Zika RNA genomes. We then perform library preparation following the SHAPE-MaP protocol to generate cDNA libraries compatible for Illumina sequencing.

### NAI-MaP structure probing of Hela cells

Hela cells were grown in DMEM high glucose media (Thermo Fisher Scientific), supplemented with 10% FBS, 1% Pen-Step to 70-80% confluency. Cells from 1 10 cm plate was trypsinized, washed once with PBS, and resuspended in 475 μl of PBS. 25 μl of 1M NAI was added to the cells at 37°C for 15 min. Total cellular RNA was extracted using TRIzol extraction, followed by passing through the RNeasy column (Qiagen). We performed structure libraries for sequencing following the SHAPE-MaP protocol.

### Pair-wise interactome mapping inside virions

Freshly collected virus was treated with a 1:500 volume of 20 mM biotinylated psoralen (2 μl of biotinylated psoralen in 1 ml of virus in cellular media) and incubated at 37°C for 10 min. The treated viruses were then spread onto a 10 cm dish and UV irradiated at 365 nm for 20 min on ice to crosslink the interacting regions. The crosslinked virus genomes were then extracted using TRIzol LS reagent (Thermo Fisher Scientific) following the manufacturer’s instructions. We performed SPLASH libraries similarly to the published protocol in Aw et al^18, 25^.

### Pair-wise interactome mapping of dengue and Zika viruses inside human cells

Huh7 and human neuronal precursor cells were infected with dengue 1 and Zika viruses at an MOI=1 for 30 hours. The cells were washed twice with PBS. The cells were then incubated with 200μM biotinylated psoralen and 0.01% digitonin in PBS for 10 min at 37 °C, and irradiated at 365nm of UV on ice for 20 min^25^. Total RNA was then extracted from the cells using TRIzol reagent according to manufacturer’s instructions. We performed SPLASH libraries similarly to the published protocol in Aw et al^18, 25^.

Translation inhibition was carried out by treating dengue1 infected Huh7 cells (30 hours post infection, MOI=1) with a final concentration of 2 μg/ml harringtonine for 5 min at 37 °C. 200μM biotinylated psoralen and 0.01% digitonin in PBS were then added to the cells for another 5 min. The cells were irradiated at 365nm of UV on ice for 20 min to crosslink the pair-wise interactions^25^, and total RNA was extracted using TRIzol. We performed SPLASH libraries similarly to the published protocol in Aw et al^18, 25^.

### Ribosome profiling of dengue1 virus in Huh7 cells

Huh7 cells were seeded in DMEM high glucose media (Thermo Fisher Scientific), supplemented with 10% FBS, 1% Pen-Step to 50-70% confluency. The cells were infected with dengue1 virus (MOI=1) in serum free media for 1 hour and grown in DMEM high glucose media (10% FBS, 1% Pen-Step) for 20 hours. To harvest the polysomes, we added 1:1000 volume of 100 μg/ml of cyclohexamide to the cell media at 37°C for 10 min. We then typsinized the cells (trypsin contains 0.1 μg/ml of cyclohexamide), washed them with PBS (containing 0.1 μg/ml of cyclohexamide) and lysed the cells using mammalian lysis buffer from the TruSeq Ribo Profile kit (Illumina). We prepared the ribosome profiling library using the TruSeq Ribo Profile kit (Illumina) following the manufacturer’s instructions.

### Functional assay to test for virus fitness

The entire genomes of dengue1 and Zika French Polynesia viruses were cloned into five overlapping fragments. Mutagenesis was performed on individual fragments and the genome was then assembled using Gibson assembly (NEBuilder HiFi DNA Assembly Master Mix) by mixing 0.1 pmole of all five fragments together. The Gibson assembled fragments are then transfected into HEK293T cells using Lipofectamine 2000 (Thermo Fisher Scientific). We collected the media from the HEK293T cells after 72 hours and infected C6/36 cells to amplify the virus. We collected the supernatant from C6/36 cells after five days post-infection and performed plaque assay to determine the virus titres. We then normalized all the wildtype and mutant viruses and infected Huh7 cells at an MOI=1 and MOI=0.1. Both the supernatant and the cellular RNAs were harvested after 24 and 48 hours and the amount of virus inside the cells and released into the supernatant was determined by qPCR. To double check that the mutant virus sequence is indeed what we engineered it to be, we performed RT-PCR on dengue and Zika genomes, using RNA extracted from infected Huh7 cells 48 hours post-infected. We then performed sanger sequencing of the virus genome to confirm the sequence of the mutant.

## Data analysis

### Shape-MaP Analysis

After quality filtering with a Phred cutoff of 25, reads were aligned against the published reference genomes (Table S1) for DENV serotypes 1-4 and the four ZIKV strains using Bowtie *2* at the highest default sensitivity setting ^26^. These alignments were performed separately for the data sets from experiments treated with DMSO and NAI respectively, as well as the denatured experiment. Mutations were counted separately in each set and subsequently expressed as a shape score by calculating (M_NAI_-M_DMSO_)/M_denat_ at each position as outlined by Weeks and co-workers^12^. Subsequently, shape scores discretized for each nucleotide position by classifying every score below 0.5 as low reactivity and at or above 0.5 as high reactivity. Discretization allowed us to easily compare structured and unstructured stretches across different viruses as well as to characterize the length and distribution of structured and unstructured motifs along the genomes. We calculated the distribution of continuous segment lengths and segment lengths with two transitions as would be expected for hairpin or helices containing a bulge. We compared these metrics with a variety of other types of RNA such as Xist mRNA, 28S rRNA and random controls^27^. Significance levels in the difference of distributions was calculated using the Wilcoxon rank-sum test.

### Sequence analysis of different DENV and ZIKV genomes

We performed multiple sequence alignment of the reference genomes using the program MAFFT with the *–genafpair* option ^28^ to align the various virus strains and species to one another. The reference alignment is provided as an supplementary material. Subsequently, a multiple sequence alignment (MSA) score was calculated for each position by determining highest number of identical pairs among all sequences and dividing it by the number of possible pairs at that position, i.e. if 7 out of 8 sequences are identical at position 1234, then the MSA score would be calculated as 21(number of identical pairs in the 8 sequences)/28(number of unique pairs in 8 sequences)=0.75. We then analyzed the degree of sequence conservation at locations classified as high-reactivity (>0.5) vs. low-reactivity (<0.5) according to NAI-MaP reactivity.

#### Analysis of covariation within viral species

We retrieved all full genome sequences for dengue 1-4 and zika from the Virus Pathogen Resource (VIPR, https://www.viprbrc.org), resulting in 1850 (DEN1), 1420 (DEN2), 923 (DEN3), 220 (DEN4), and 540 (ZIKV) full genome sequences respectively. After aligning the sequences for each viral species separately using mafft with the -localpair’ option, we compared all pairs of sequences and counted all covariations at each position. This yields comprehensive covariation information across DEN1-3, while the abundance of covariation data for DEN4 and ZIKV is somewhat limited. For DEN4, the small number of available full genome sequences imposes a detection limit on covariations, whereas for ZIKV, the set of full genome sequences contains a high number of recent, and similar sequences mostly originating in the South American outbreak, which exhibit narrow variability and hence provide only limited information on covariation. Across the manuscript, evidence of covariation is indicated as green circles for the respective base pairs.

### Analysis of sequence conservation at structurally similar or different bases in DENV and ZIKV genome

Based on shape reactivity, bases were classified as high (H) or low (L) using a cut-off of 0.5. The cut-off for NAI-Shape-MaP was optimized to have the best discriminative power for the known 28S structure. To assess whether structures are conserved across DENV and ZIKV viruses, majority votes were applied at each position of the reference alignment. Bases whereby the four DENV subtypes or the four ZIKV strains have either 3/4 or 4/4 identical shape classes (e.g. HHHL would be classed as H, LLHL would be classed as L) as classified as structurally similar. Classes without majority were identified as indeterminate (e.g. HLHL or HHLL) and were not included in the subsequent analysis. Subsequently, we divided the genome into positions that have identical majorities for DENV and ZIKV (same, S), and positions where the majority structure is different between DENV and ZIKV (different, D). We then calculated the distribution of multiple sequence alignment scores within each class. Significance levels in the difference of distributions was again assessed using the Wilcoxon rank-sum test.

### Structure modelling of DENV and ZIKV genomes

Using the Shape-MaP data in conjunction with RNA structure prediction algorithms allowed us to create structural models for the full-length Dengue and Zika genomes. We utilized the program RNAstructure^29^ and incorporated the Shape-MaP results as additional restraints using a slope of 0.5 and an intercept of 0.0 kcal/mol for the Shape-MaP potentials. These values were obtained by optimization of structure prediction for 28S human ribosome using NAI-Shape-MaP data, otherwise using default settings.

### SPLASH long-range interactions

#### SPLASH long-range interactions

SPLASH reads were aligned to their respective reference genome using the program Bowtie 2 to perform a paired read alignment^26^. A matrix containing a count of all pairwise T positions observed in the reads was compiled. To filter random pairs, we constructed a baseline matrix by randomly shuffling all read pairs 100 times and subsequently subtracted this baseline matrix from the raw read matrix. We then considered all sites that exceed a threshold value of 10% of the maximal observed value as valid SPLASH interactions. The shuffling procedure and threshold values were optimized for the known human and E. coli ribosome structures, and allowed us to markedly improve specificity and the positive predictive value for SPLASH interactions. Long-range interactions were considered conserved if they occur in at least two strains. Plotting the precise read depth across the genome allows for identification of distinct peaks mostly within <50nt segments and thus allows the localization of the interaction sites. We analyzed the promiscuity of these long-range interactions by counting the number of alternate interaction partners we find at each site originating long-range interactions. We identified several sites that show multiple coinciding interactions for the same sequence stretch. Competing interactions were modelled using *RNAcofold* from the ViennaRNA package and visualized including the Shape-MaP and covariation data^19^.

### Peak calling of SPLASH interaction sites

The genome of each of the eight viruses were divided into non-overlapping bins of 100 bases. Each chimeric read was split and mapped to two paired bins. Paired bins that contain at least 20 chimeric reads were then selected for peak calling. The density curves of read coverage were fit (using the “density” function of R) for the left and right bins respectively. The peak of the interaction was determined to be the position with the maximum coverage density.

### Identifying regions of functional interest

Regions of functional interest were identified if they fulfill at least 2 out of 3 of the following criteria: (i) Structure similarity is in the top 30% among all regions; (ii) structured stretch length is in the top 30% of all regions; and (iii) the region is in the bottom 30% for synonymous mutation rate, indicating selection pressure on the RNA itself for both dengue and Zika. Structural similarity was assessed by calculating the local identity of the discretized shape scores. The length of structured stretches was calculated from the same data set. The synonymous mutation rate was calculated from an alignment of all dengue and Zika full genome sequences available at the time of writing. Structure models for all identified sequences including available covariation data are presented in the manuscript.

## Acknowledgements

We thank members of the Wan lab, Sessions lab, Ooi lab for discussions. YW is supported by funding from A*STAR and Society in Science - Branco Weiss Fellowship. PJB acknowledges support by funding from the Ministry of Education Singapore - MOE AcRF Tier 3 Grant Number MOE2012-T3-1-008. RGH acknowledges support by funding from the A*STAR Biomedical Research Council - Young Investigator Grant BMRC YIG 1610651031.

## Author contributions

Y. Wan conceived the project. Y. Wan, EE. Ooi, OM. Sessions, PF. Sessions designed the experiments. X.N. Lim, W.C. Ng, M.J Choy, KL. Boon, J.G Aw performed the experiments. A. Sim, R. Huber, Y. Shen, A. Wilm and P. Bond performed the computational analysis. Y. Wan organized and wrote the paper with A. Sim, R. Huber, OM. Sessions, PF. Sessions, M. Hibberd, N. Nagarajan and all other authors.

## Extended data legends

**Extended data 1. NAI-MaP identifies single stranded regions in vivo. a**, NAI-MaP reactivities on the 28S rRNA in vivo. Highly reactive NAI-MaP signals (red) map to single-stranded regions along the known secondary structure.

**Extended data 2. Structure probing of full length, dengue and Zika genomes inside virus particles. a,b**, Gel images showing that structure probing was performed on full length, intact, dengue (A) and Zika (B) genomes. Extracted RNA is run on 0.6% agarose gels using DNA and RNA ladders as reference. L is 1kb plus DNA ladder (GeneRuler), RNA L is ssRNA ladder (NEB). **c**, In vitro structure probing of the dengue1 genome using enzymatic digestion followed by deep sequencing (PARS, red line), or NAI-MaP (black line). High reactivity in NAI-MaP indicates that a base is single-stranded while high signal for PARS indicates that a base is double stranded. PARS and NAI-MaP signals are largely inverse of each other, suggesting that they are identifying double and single stranded bases at similar positions. **d**, PARS and NAI-MaP are mapped to the known 5’UTR structure in dengue1.

**Extended data 3. Analysis of NAI-MaP data in vivo. a**, Sequence identities between dengue and Zika strains. **b**, Choice of cutoff for discriminating structured and unstructured regions with NAI-ShapeMaP. Shape reactivity for 28S rRNA was split into structured and unstructured bases according to the experimentally determined structure. The dashed lines show bases classified as structured by a certain cutoff as a fraction of all bases of the true structured (black) and true unstructured (grey) class. The red curve is the difference between the black and grey curves and thus shows the ability of a certain cut-off to discriminate between structured and unstructured bases. This ability is at its highest at a cutoff of about 0.5. **c**, Violin plots showing the distribution of sequence similarity across eight viruses in positions that are structured (black) versus positions that are not structured (red). Single stranded regions tend to be more highly sequence conserved across the eight viruses as compared to double stranded regions. **d**, Violin plots showing the distribution of sequence similarity across the eight dengue and Zika viruses in positions that are structurally similar (left) and structurally different (right) across the viruses. Bases that share similarities in structure tend to share similarity in sequence.

**Extended data 4. Analysis of conserved local structural regions in dengue and Zika. a, b**, Plots showing the extent of structural similarity, double-strandedness (structured stretches), and the extent of synonymous mutation rate across the four dengue (**a**) and four Zika (**b**) genomes. Structure similarity is calculated using pearson correlation of 100 base windows along the genome. Structured structures is the inverse of NAI-MaP reactivities (The lower the NAI-MaP reactivity in the 100 base window, the more structured that window is). The top 30 percentile of each of these plots are shown as blocks in Figure 2a, and Extended data 4c. **c**, Plots show 100 nucleotide regions across four dengue genomes that have highly similar structures (grey), are highly double-stranded (black), and accumulate low levels of synonymous mutations (blue). Regions that fulfil two out of three of the above criteria are selected as consensus regions (purple) and are potentially functionally important. **d**, Violin plot showing the distribution of Shannon entropies in the conserved 16 dengue structures (selected regions) versus all regions along the genome. The conserved structures show lower Shannon entropies, suggesting that they form unique structures. *P*-value was calculated by Wilcoxon Rank Sum test.

**Extended data 5. Incorporating NAI-MaP into structure modelling to achieve accurate structure models. a**, Table showing the percentage of correct base pairs that are modelled in 16S and 23S rRNA with and without incorporating NAI-MaP reactivities into the RNAstructure program. **b**, Violin plot showing the distribution of co-varied bases found within NAI-MaP constrained structure models versus random. *P*-value is calculated by Wilcoxon Rank Sum test. **c,d**, Histogram of the distribution of the length of stems that are present in modelled dengue and Zika RNA structures after incorporating NAI-MaP reactivities into RNA structure program. **e**, Table containing information on the length of helixes in dengue and Zika serotypes.

**Extended data 6. Structure models of conserved Zika regions**. Structure models of the Zika French Polynesia genome using NAI-MaP as experimental constraints, for the 12 consensus regions identified in Extended data 4c. The NAI-MaP reactivities (red, orange, black, indicating single, intermediate and double stranded regions respectively) and covariation information (green circles) are mapped onto the structures.

**Extended data 7. Ribosome profiling along the dengue1 genome in Huh7 cells. a**, Fourier transform of the ribosome footprints along the dengue1 genome shows a three nucleotide periodicity. **b**, Ribosome footprints normalized by mRNA count along the dengue1 genome. Base1 refers to A in the AUG start codon. The red bars indicate the footprints present in the coding sequence and the black bars indicate footprints in the 5’UTR sequence. **c**, Boxplot showing the distribution of NAI-MaP reactivity at the strongest 1% and 10% of ribosome profiling pause sites, versus all sites. **d**, The ribosome pause site is located in segment 12 consensus structure region and in a region of low Shannon entropy, suggesting that it is likely to be present in a defined structural region. **e**, Table showing the average distribution of codon frequencies in 2, 4, 6, 8, 12, 18 codons (local mean) around the GAU codon at the 7272 ribosome pause site. The global mean indicates the average codon frequencies for all codons in the dengue1 genome. Codon frequencies surrounding the ribosome pause site are not significantly different from codon frequencies along the entire dengue1 genome, suggesting that codon usage is unlikely to be the main reason behind the strong ribosome pause site.

**Extended data 8. New long-range interactions are identified in dengue and Zika viruses. a**, 2D matrixes showing the location of pair-wise RNA interactions along the dengue1 genome inside virions (left), upon random shuffling (middle), and upon filtering against random shuffled interactions (right). **b**, Top: arc plots showing one set of alternative interactions found along the Zika French Polynesia genome. Bottom: structure models of the pair-wise interactions for the alternative structures, using the program RNAcofold. The NAI-MaP reactivities (red, orange, black, indicating single, intermediate and double stranded regions respectively) and covariation information (green circles) are mapped onto the structures. **c**, Schematic of the functional assay to test for mutant virus fitness (Methods).

**Extended data 9. Dengue and Zika can use different but nearby sequences to achieve genome organization. a**, Arc plots showing conserved long-range interactions that are present in at least 2 dengue genomes in virions. A long range interaction that brings bases 4000 into close proximity to bases 9000 is seen in both dengue and Zika genomes (blue). **b**, Structure models of the predicted interactions between bases 4000:9000 in the different dengue and Zika viruses are generated using the program RNAcofold. NAI-MaP reactivities (red, orange, black, indicating single, intermediate and double stranded regions respectively) and covariation information (green circles) are mapped onto the structures.

**Extended data 10. Long-range interactions in dengue virions are disrupted inside host cells. a**, 2D matrixes showing the location of pair-wise RNA interactions along the dengue1 genome inside cells (left), upon random shuffling (middle), and upon filtering against random shuffled interactions (right). In cell pair-wise interactions are enriched for short, local interactions along the diagonal of the plot. **b**, Arc plots showing long-range interactions in dengue 1 virus that are present inside virion particles (top) versus inside host cells (bottom).

